# RNA G-Quadruplex Emerges from a Coil-Like Ensemble via Multiple Pathways

**DOI:** 10.1101/2025.01.07.631675

**Authors:** Pavlína Pokorná, Vojtěch Mlýnský, Jiří Šponer, Petr Stadlbauer

## Abstract

RNA G-quadruplexes (rG4s) are vital structural elements in gene regulation and genome stability. In untranslated regions of mRNAs, rG4s influence translation efficiency and mRNA localization. Additionally, rG4s of long non-coding RNAs and telomeric RNA play roles in RNA processing and cellular aging. Despite their significance, atomic-level folding mechanisms of rG4s remain poorly understood due to their complexity. We used enhanced-sampling all-atom molecular dynamics simulations to model the folding of an r(GGGA)_3_GGG sequence into a parallel-stranded rG4. The folding pathways suggest that RNA initially adopts a compact coil-like ensemble, marked by dynamic guanine stacking and pairing. The three-quartet rG4 builds up gradually from the coil via diverse routes involving strand rearrangements and guanine incorporations. While the folding mechanism is multi-pathway, various two-quartet rG4s seem to be a common transitory ensemble for most routes. Our simulations also exposed force-field imbalances, with the predicted folding free energy of +12.5 kcal/mol deviating from experiments. Additionally, the enhanced sampling protocol, combining well-tempered metadynamics with solute tempering, faced productivity challenges on the multidimensional free-energy surface. Overall, this study provides atomistic insights into rG4 folding, emphasizing compact coil-like ensembles as key precursors, while revealing limitations in simulating non-canonical RNA structures.

## Introduction

Guanine quadruplexes (G4) are important non-canonical nucleic acid structures formed by both DNA and RNA molecules.^1^ Guanine-rich sequences capable of G4 formation are found across all domains of life.^2^ Whilst the list of biological roles of DNA G4s (dG4) has been expanding for more than two decades, the existence of RNA G4s (rG4) *in vivo* has been demonstrated rather recently.^3, 4^ Similarly to dG4, rG4s have been suggested to play major regulatory roles in gene expression^5–9^ and at maintaining genome integrity at telomeres.^10–13^ As such, rG4s present a promising target for treating various diseases, including cancer. Besides their biological roles, G4s are also used as building blocks in nanomaterials or utilized in biosensing.^14, 15^

G4s are composed of at least two stacked guanine quartets. Each quartet is formed by four guanines, which are H-bonded in a cyclical fashion using their Watson-Crick and Hoogsteen edges for pairing (Figure 1A).^16^ In the middle of the quartet, there is a cavity, which upon quartet-quartet stacking, forms a channel that runs through the G4 stem and binds cations stabilizing the G4 structure.

**Figure 1.**
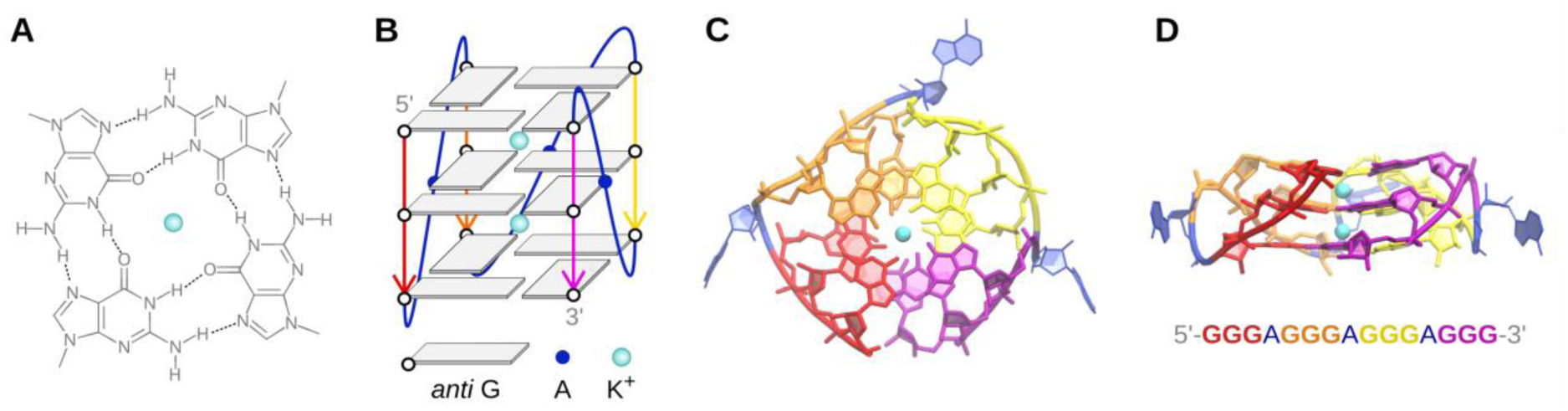
RNA G-quadruplex. (A) Guanine tetrad; the sugar moiety is not shown for clarity. (B) Sketch of the parallel-stranded G4 (GGGA)_3_GGG; backbone of the first, second, third, and fourth G-tract (G- column, G-strand) is colored in red, orange, yellow and mauve, respectively, the loops are blue. (C) Top view and (D) side view of the actual rG4 structure used as a reference in this work; G-tracts are colored as in (B), hydrogens are not shown for clarity.

There are some interesting differences between dG4 and rG4; dG4 can adopt diverse topologies differing in how the backbone winds around the quartets, leading to variability of G- strand orientations (parallel or antiparallel) and loop types (propeller, lateral, diagonal or V- shaped).^17, 18^ Different dG4 topologies are intimately interrelated with specific *syn*/*anti* patterns of the glysosidic angles in the G-strands. Importantly, riboguanosine adopts the *syn* conformation much less likely than deoxyriboguanosine. As a result, rG4s adopt basically exclusively the all-*anti* parallel-stranded topology with propeller loops (Figure 1).^19^ .^20^

The folding kinetics of various G4s has attracted considerable attention. Most of the effort has been paid to DNA sequences, mainly the human telomeric sequence d(GGGTTA)_3_GGG and its variants, whereas rG4 folding has been understudied. The folding process (its kinetics, involved intermediates, and its dependence on sequence, starting state and environment) of G4s is not only an interesting physical-chemistry phenomena *per se*, but it may also affect the biochemical roles of G4-forming sequences.^20–22^

In general, dG4 folding kinetics (i.e., reaching thermodynamic equilibrium in a specific folding experiment) may be very slow; folding times from minutes up to weeks have been reported for dG4.^20–25^ The long folding timescales observed in many dG4 experiments can be explained by so called kinetic partitioning,^26^ i.e., presence of two or multiple long-living states (deep free-energy basins) on the G4 free-energy landscape which are acting with respect to each other as off-pathway kinetic traps. It has been argued that the key long-living states are diverse dG4 folds with different *syn*-*anti* patterns in their G-tracts.^27–31^

In this context, the rG4 folding landscape is quite different. It has been theorized^27, 32^ and subsequently demonstrated^33^ that rG4 folding is faster compared to its dG4 counterparts. The reason is that riboguanosine is much less likely to adopt the *syn* conformation. This generally eliminates the competing long-living G4 folds with non-native *syn*-*anti* G conformation patterns from the free-energy landscape and in addition increases the chance that upon mutual encounter the guanines will be oriented in a manner favorable for a successful rG4 formation.^32^ Thus, the folding landscape of an rG4 is much less rugged than that of dG4.

Unfortunately, G4 folding is difficult to monitor at fine resolution using experimental methods. Techniques, such as optical/magnetic tweezers usually combined with FRET,^21, 30, 31, 34–43^ CD,^24, 25, 42, 44, 45^ NMR^22, 28, 33, 46, 47^ or mass spectrometry^29, 48, 49^ have been used to study the folding process, but none of them has sufficient temporal and spatial resolution to provide atomistic details of the folding process; most direct information about the folding can be obtained by time-resolved NMR experiments.^22, 28, 33, 46, 47^ Therefore, structural interpretations of the primary experimental data are only indirect, often based on intuition and presumptions. Computer simulations have been used to provide the atomistic aspect of the G4 folding.^32, 38, 44, 50–73^ Standard (unbiased) all-atom explicit-solvent molecular dynamics (MD) simulations are the most accurate tool, but they are severely limited by the affordable time scale, reaching up to hundreds of microseconds, which is not enough to capture the entire G4 folding process.^74^ Thus, standard MD simulations have been used to analyze properties of diverse types of structures that may participate in the G4 folding process and have indirectly suggested plausible mechanisms of the folding, including the kinetic partitioning.^32, 50, 51, 53, 60, 61, 63–65, 69, 71^ Besides that, many atomistic simulation studies used diverse enhanced-sampling methods to probe the G4 folding.^32, 38, 53, 56–59, 61, 63, 65–68, 70, 72, 73, 75, 76^ These methods are designed to investigate broader portions of the free-energy landscape. However, these approaches can also distort the folding mechanism, mainly due to excessive dimensionality reduction associated with the use of so-called collective variables (CVs) along which the sampling is enhanced (for a review see ^27, 74^). Yet another approximate option is to use non-atomistic (coarse-grained, CG) methods.^53, 55, 62, 77–80^ Accuracy of all simulation methods obviously depends on the quality of the potential energy function (the force field) describing the nucleic acid molecule.^81^

All-atom MD simulations have also been used to study putative rG4 folding intermediates, namely hairpins formed by the sequence rGGGAGGG and telomeric repeat-containing RNA (TERRA) fragment rGGGUUAGGG.^32^ Although these sequences are theoretically capable of forming G-hairpins, the simulations predicted that “G4-like” or “ideal” G-hairpins are unstable in the parallel orientation. Instead, a cross-like state was more favored. However, unfolded, coiled and antiparallel hairpins were even more frequently populated. The key distinction between the “G4-like” and cross-like states lies in their pairing patterns (Supporting Figure S1): in the “G4-like” state, all Gs are Hoogsteen-paired with their “correct” partners – matching those in a fully folded G4 structure. In contrast, the cross-like state represents an ensemble of structures where the strands are rotated relative to one another. This rotation results in Gs being paired in a less specific manner, with some Gs possibly remaining unpaired. The simulations have also suggested that parallel G-triplexes are likely rather unimportant intermediates due to their low structural stability. Instead, compact but rather unstructured coil-like states have been suggested as potential seeds from which the G4 fold can emerge. CG simulations using the three-interaction-site model in implicit solvent^82, 83^ investigated the folding of the whole r(GGGA)_3_GGG sequence.^77^ They have suggested that the rG4 folding is a multi-pathway process, with salient formation of stacked rGGG columns. Unlike the unbiased all-atom simulations, the CG model predicted that the G-tracts straightforwardly interact to form well-structured parallel G-hairpins. Then rG4 arises either by consecutive attachment of two additional G-columns via a parallel G-triplex, or by merger of a pair of G-hairpins. No coil-like and cross-like structures were reported. The struggle to capture the rG4 folding by computational tools and the complexity of the rG4 free-energy landscape can be demonstrated by recent simulations using the SimRNA CG model, in which the RNA chain folded into the native rG4 only when the native G:G base pairs were predefined manually.^84^

We have recently reported folding simulations of d(GGGA)_3_GGG dG4 using enhanced-sampling MD, combining replica-exchange approach with metadynamics.^66^ The used sequence has the minimal loop length (one nucleotide) and thus folds just into a single dominant topology, the parallel-stranded all-*anti* dG4. The study highlighted the multi-pathway nature of the folding process devoid of distinct well-defined simple intermediates. The simulations suggested that the molecule first forms a loosely structurally defined compacted coil-like ensemble, from which the dG4 structure itself emerges through multiple small consecutive steps. At the same time, however, the simulations predicted that the folded dG4 does not correspond to the global minimum on the free-energy landscape. This has been attributed to inaccuracy of the simulation force field, which does not capture the proper balance between the unfolded and folded ensembles; the correctly folded structure has nevertheless been reached with the help of the used CV and the associated biasing potential. In the present work we use the same method to probe the free energy landscape of r(GGGA)_3_GGG rG4. The results enable us to assess similarities and differences in the folding processes of DNA and RNA G4s. The data suggest that the folding of parallel-stranded rG4 is in many aspects similar to folding of the analogous dG4 with single nucleotide loops; it may proceed via various intermediates, including cross-like triplexes, two-layered G-triplexes and slip-stranded rG4. Importantly, all of them are again emerging from the loosely defined coil-like ensemble, which thus serves as the key transitory ensemble of the folding process.

## Methods

### Starting structure

The folding simulations were initiated from a fully extended r(GGGA)_3_GGG RNA strand, built by nucleic acid builder in the A-RNA conformation.^85^ The RNA was described with OL3 all-atom force field^86–88^ and solvated in explicit SPC/E^89^ water under the presence of 0.15 M excess KCl.^90^ The minimal distance between the extended RNA strand and the solvent box border was 25 Å.

To create the reference and target rG4 structure, we took the parallel-stranded dG4 2LEE,^91^ mutated the cytosines in the single-nucleotide loops to adenines and converted the whole molecule into RNA. The molecule was then carefully equilibrated to assure that it adopts a proper rG4 structure which can be used for evaluation of the simulated structures by the εRMSD metric (see below).^92^

### Simulation protocol

We used the replica-exchange solute-tempering simulation protocol (REST2)^93^ coupled with well-tempered metadynamics^94^ (thus called ST-metaD) to enhance the sampling. In total, we ran four ST-metaD simulations, each with 16 replicas (Table 1). The replicas spanned the effective temperature ladder from 285 to 500 K and the metadynamics bias potential was built in each replica. As a collective variable (CV), we used the inter-tract εRMSD metric,^66, 92^ which describes the fit of guanine base positions and orientations with respect to the reference folded rG4 structure. In general, εRMSD is a metric tailored for a description of nucleobase interactions in nucleic acids. It is sensitive to changes in base-pairing and stacking and thus is suitable for monitoring of the G4 folding process. Inter-tract εRMSD is a modification that considers only εRMSD contribution between G-tracts, while ignoring stacking of guanines within a single G-tract to avoid an excessive promotion of intra-tract guanine stacking at the expense of inter-tract encounters (Supporting Figure S2).^92^

**Table 1.** List of rG4 folding simulations.

| Simulation number | Replicas x simulation time (μs) | $\Delta G_{\text{fold}}$ (kcal/mol)<sup>a</sup> | Number of folding events |
| --- | --- | --- | --- |
| 1 | 16 x 4.0 | 9.6 | 4 |
| 2 | 16 x 4.0 | 15.6 | 3 |
| 3 | 16 x 4.0 | 12.3 | 3 |
| 4 | 16 x 2.0 | - | 1 |

<sup>a</sup> Snapshots with εRMSD to the reference folded structures smaller than 0.7 were considered as part of the folded ensemble (see Supporting Figure S3) while the remaining snapshots were counted as the unfolded ensemble.

While we only used the reference replica (with an effective temperature of 298 K) to calculate ΔG_fold_, we have monitored the development in the continuous (demuxed) trajectories to visualize the folding events. As folding events we consider portions of the continuous trajectories, so-called reactive trajectories, which bring the simulated molecule from a clearly unfolded state to a fully folded G4. For the reactive trajectories we monitor in detail the development of simulated structure to capture the folding pathway. Obviously, the folding pathways are necessarily affected by the used enhanced-sampling protocol, i.e., by travel of the continuous replicas through the REST2 replica ladder and mainly by the used biasing CV. We still suggest that the simulations reflect important features of the folding landscape.^95^

Before the ST-metaD simulations, the systems were relaxed to avoid unfavorable contacts and voids. The structures were minimized and equilibrated in a series of steps with gradual decrease of position restraints. Then we performed a 500 ns long nVT simulation, from which we extracted 16 starting structures for the 16 replicas of the actual ST-metaD simulation to heterogenize the sampling. We used the V-rescale thermostat to keep the temperature at 298 K.^96^ We employed the SHAKE algorithm^97^ together with the hydrogen-mass repartitioning,^98^ which allowed us to use a 4-fs integration time step. Simulations protocols are available in Supporting Information.

While the bias potential in ST-metaD simulations converges after ∼2-3 microseconds in our simulations, it is certainly not possible to obtain complete sampling of the folding landscape due to the very rich conformational space of the simulated molecule. Therefore, while the ST-metaD protocol combined with εRMSD is, to our knowledge, one of the most powerful methods to simulate G4 folding, it could only sample lower singles of folding events in 16-replica simulations. Out of the four ST-metaD simulations we carried out, one sampled only a single folding event after 2 microseconds and predicted a very high ΔG_fold_. Thus we stopped that run. At the same time, we extended the other three runs to 4 microseconds. Detailed analyses of the individual continuous trajectories (εRMSD and travel through the replica space) are provided in the Supporting Information. Limitations of the ST-metaD method that we encountered in course of this study are discussed in the Results and Discussion section.

## Results and Discussion

We have performed four simulations of the RNA G-quadruplex (rG4) forming sequence r(GGGA)_3_GGG and observed eleven rG4 folding events (Tables 1 and 2, Supporting Movies SM1-SM11). Most importantly, the simulations show that the folding proceeds via a loosely defined coil-like ensemble, from which rG4 gradually arises by following diverse micro-routes and intermediates. High ΔG_fold_ energy of rG4 nevertheless indicates that the force field is unable to reproduce the rG4 thermodynamic stability correctly; the folding is achieved because it is boosted by the bias potential along the used CV.

**Table 2.** List of folding events.

| Reactive trajectory number<br>(as in Figures 3, 4, S6, and S7)<sup>a</sup> | Simulation number | Demuxed trajectory number<br>(as in Figures S3 and S4) | Folding time based on εRMSD (ns)<sup>b</sup> | Time window presented in Figure 3 and Supporting Movies (ns) |
| --- | --- | --- | --- | --- |
| 1_1 | 1 | 8 | 1444 | 1000-1500 |
| 1_2 | 1 | 10 | 946 | 0-1000 |
| 1_3 | 1 | 14 | 70 | 0-100 |
| 1_4 | 1 | 7 | 232 | 0-250 |
| 2_1 | 2 | 5 | 205 | 0-250 |
| 2_2 | 2 | 6 | 2739 | 1500-2900 |
| 2_3 | 2 | 2 | 75 | 0-100 |
| 3_1 | 3 | 4 | 31 | 0-100 |
| 3_2 | 3 | 10 | 1559 | 0-1600 |
| 3_3 | 3 | 11 | 127 | 0-150 |
| 4_1 | 4 | 11 | 64 | 0-100 |

<sup>a</sup> Reactive trajectory is a part of continuous (demultiplexed, demuxed) trajectory that contains the series of structural transitions events (the shortest path) leading from the fully unfolded state to the fully folded rG4. By the fully folded rG4 we mean a structure with εRMSD to the reference folded rG4 structure smaller than 0.7. The fully unfolded state is not precisely defined but was identified through visual inspection of graphs tracking individual structural features of the simulated molecule. It corresponds to a point where the molecule lacks any discernible rG4-like characteristics, as illustrated in Figure 3. “Continuous” means that we follow the trajectory of a given (initially fully unfolded) molecule as it travels across the replica ladder, instead of monitoring the structures as they populate the discontinuous replica at a given temperature.
<sup>b</sup> Time at which the specific folding event was completed in given simulation run. See detailed timelines in Supporting Figure S3.

### Folding free energy

The simulations predict that the rG4 ΔG_fold_ energy is about +12.5 kcal/mol on average (Table 1), with the fully-folded rG4 separated from the rest of the ensemble by a relatively low energy barrier (Figure 2). The bias potential is converged and the replica space seems sampled sufficiently well (Figure 2 and Supporting Figure S4), but the folding events were rather rare. In comparison to the analogous dG4 studied before,^66^ rG4 ΔG_fold_ is lower than that of dG4 by 4.5 kcal/mol while the shape of the free-energy profile along the εRMSD CV is rather similar (Figure 2 and Supporting Figure S5). Thus, in both cases the global minimum ensemble is far from the folded G4, suggesting that the AMBER force field struggles to capture G4 thermodynamic stability in general.

**Figure 2.**
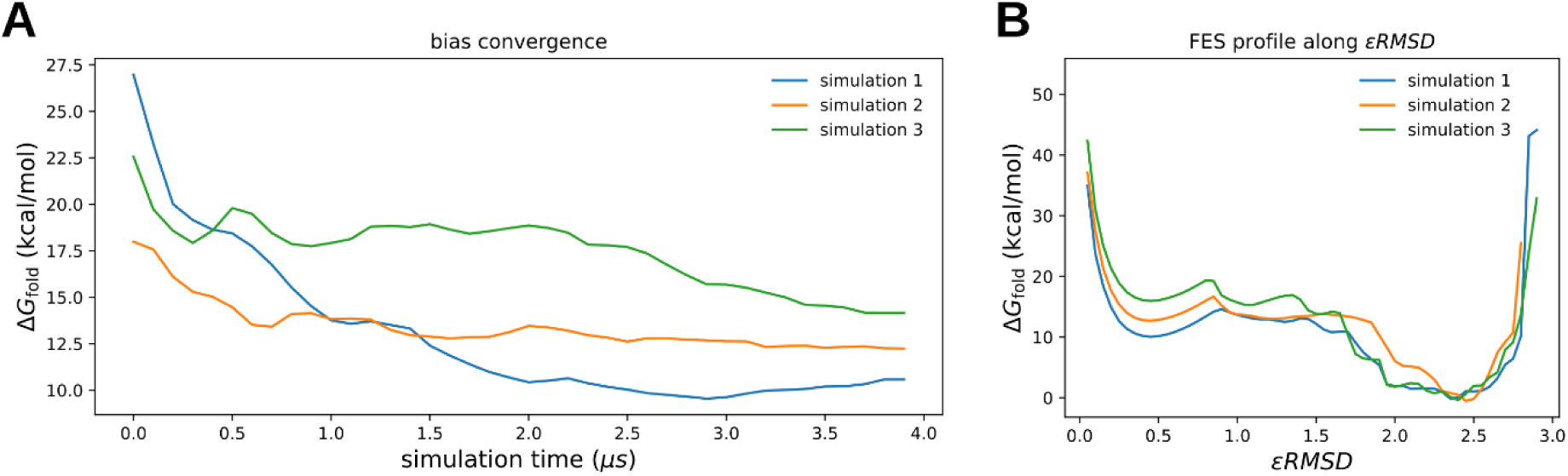
Folding free energy. (A) Bias convergence in the three 4-µs rG4 folding simulations. Structures with εRMSD from the folded reference < 0.7 were considered part of the folded ensemble. (B) ΔG_fold_ along the εRMSD CV; values are calculated using a bin width of 0.1. See Supporting Figure S5 for a comparison with DNA dG4.^66^

The evident global force-field imbalance is most likely dominantly caused by the lack of polarization in pair-additive force fields.^99^ This leads, e.g., to well-documented large-scale inaccuracies in description of the ion-ion and ion-G-tetrad interactions in the G-stems.^100^ Treatment for these inaccuracies may come from the polarizable force fields, which have already been shown to improve the behavior of ions in the G-stems, however, the description of the loop regions is still suboptimal.^101–103^ Reparametrization of non-bonded interactions specifically designed for G4 structures is another possible approach that may improve the calculated free energy of G4 folding.^57^ Besides the absence of polarization, we have also several times noticed that the propeller loops in atomistic simulations may be spuriously destabilized by the current force fields though no clear origin of such under-stabilization has so far been suggested.^27, 32, 63, 69, 104^ In summary, although our folding free-energy estimation is clearly offset by a huge margin, the simulated folding pathways should still provide some valid insights into structural ensembles populated on the G4 folding landscape, because the ST-metaD potential energy bias overcomes the force-field misbalance.

### Multi-pathway folding of rG4 from the coil-like ensemble

From the four ST-metaD (GGGA)_3_GGG RNA simulations (cumulative time of 224 µs) we obtained eleven folding events (Figures 3 and 4, Table 2, Supporting Movies SM1-SM11). Although the pathways of the individual folding events were very diverse and each was unique (Figures 3 and 4, Supporting Movies SM1-SM11), we identified a few general features that were common among the simulations. Typically, the folding started by a compaction of the extended RNA chain by formation of H-bonds between two G-tracts (e.g., the first and last (fourth) G-tract (labelled as **1-4**), or the second and third one (**2-3**)), although not necessarily leading to the correct native rG4-like pairing. Although the guanines within the individual G-tracts tended to be stacked, we commonly observed states with only two guanines stacked while the third one was then typically stacked on another G-tract or with the loop adenine. Notably, these structures with two G:G base pairs were not necessarily the ideal G4-like G-hairpins, but they often had the G-tracts rotated into the cross-like shape (Supporting Figure S1). The initial pairing led to (or was followed by) a compaction of the chain into the coil-like arrangement. This intermediate state cannot be characterized by some structure-specific interactions, as it was a structurally diverse ensemble, which contained parallel, cross-like, as well as antiparallel hairpins. The hairpins could be formed by pairing of two or three guanines either from neighboring G-tracts (i.e., G-tracts **1-2**, **2-3**, or **3-4**) or between the first and last G-tract (**1-4**). The pairing and the whole coil-like ensemble were very dynamic, so when a hairpin was formed, it could have been unfolded later. Importantly, the inclusion of ions usually happened in this coil-like state, thus preceding formation of the tetrads, and was dynamic, too (Figure 3).

**Figure 3:**
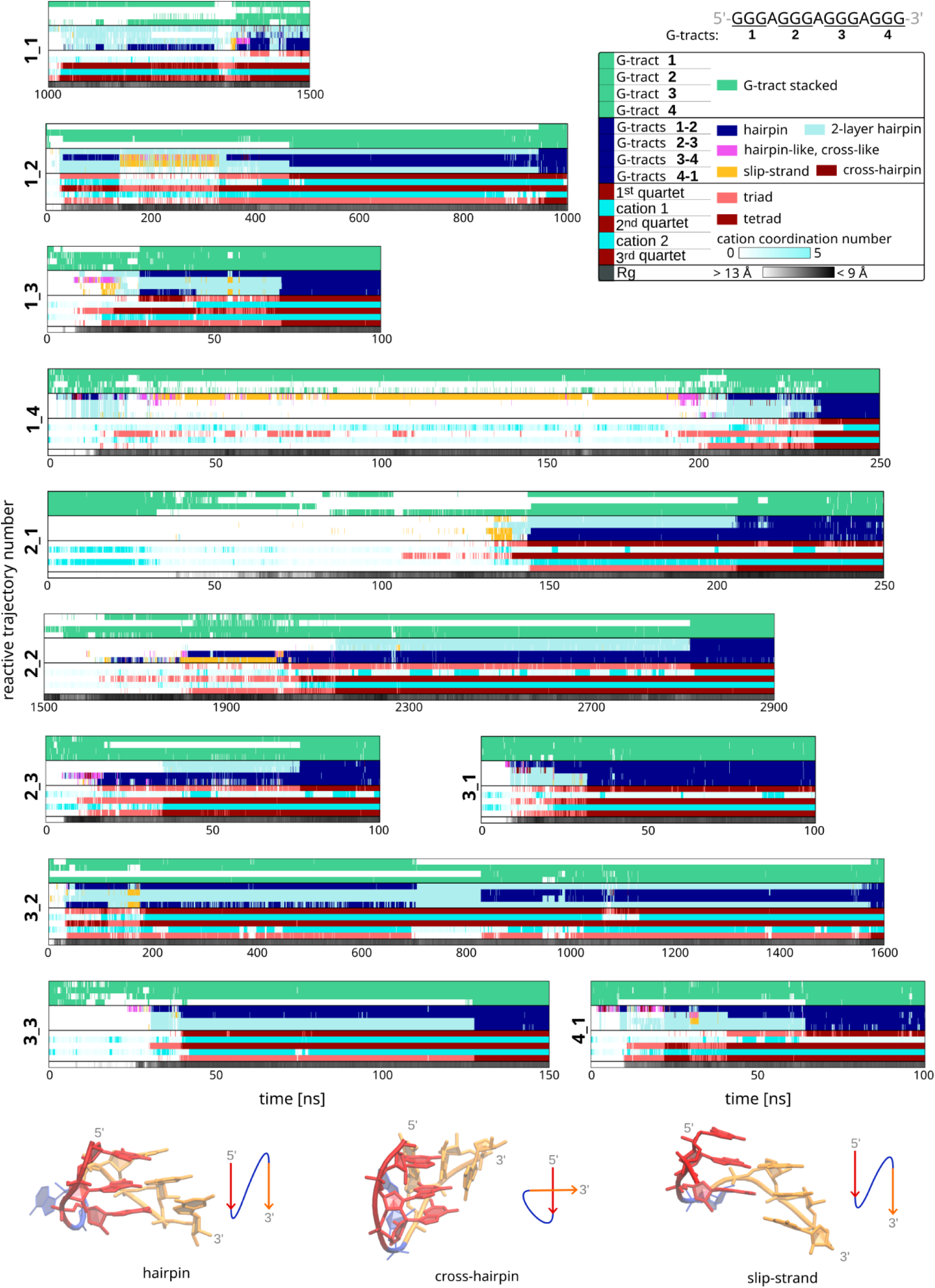
Development of key structural features in the eleven folding events, i.e., the reactive trajectories. The respective simulation runs and continuous trajectories are indicated on the y-axis (see Table 2). Irrelevant trajectory portions (not considered as part of the folding event) are omitted in the graphs where relevant. The four top-most stripes in each graph monitor stacking of the G-tracts; the green color means that all three guanines are stacked. The next four stripes monitor mutual orientations of the neighboring G-tracts (see the detailed legend to the Figure in the middle-right). The following five stripes indicate formation of the individual rG4 layers (triads or tetrads) and cation coordination between them. When all the stripes are colored as in the left column of the legend, the rG4 is fully formed. The last stripe shows the overall radius of gyration (Rg); note that Rg does not appear to be a good descriptor of the folding (see also below). Representative examples of mutual orientation of two G-tracts are shown in the bottom left corner. Actual structures from seven folding events are shown in Figure 4. Verbal description and movies of the events are provided in Supporting Information and the development of the whole trajectories is shown in Supporting Figures S6 and S7.

**Figure 4.**
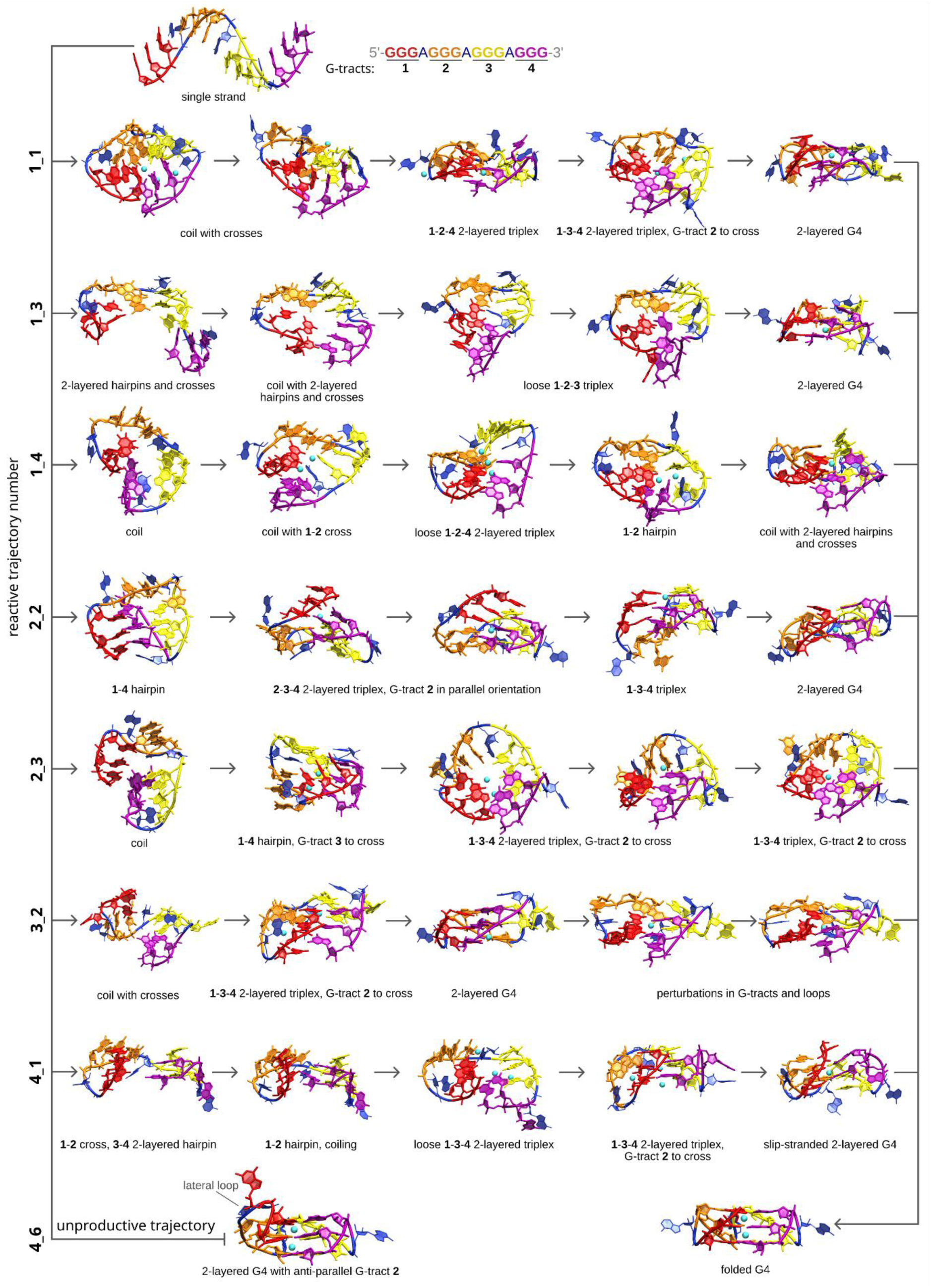
Example of seven rG4 folding pathways with various intermediates and a misfolded rG4 observed in the four ST-metaD simulations. The reactive trajectory numbering corresponds to that in Figure 3 and Table 2). A misfolded rG4, i.e., an unproductive trajectory, is shown at bottom left. The shown intermediates are snapshots from broad ensembles of similar structures. The events are also visualized in Supporting Movies.

The rG4 then gradually grew from this coil-like ensemble via numerous incremental rearrangements. We usually observed formation of two-quartet rG4 intermediate ensembles, which could be slip-stranded, i.e. with vertically shifted G-tracts (Figures 3 and 4). We did not detect any formation of states that would contain just one quartet. Complete fully folded three-quartet rG4 then emerged by incorporation (zipping) of the missing guanines into the two-quartet intermediate and strand-slippage to achieve the correct pairing, if necessary. Importantly, three-layered G-triplex with the fourth G-tract in the cross-like orientation to it was observed in two simulations, and it did not lead to rG4 formation; i.e., it was an unproductive attempt. This suggests that the pathways via 2-quartet intermediates are more important, most likely for entropic reasons. Although the simulated rG4 typically adopts the parallel all-*anti* topology, we detected formation of a misfolded two-layered rG4, which had an antiparallel G-tract with all guanines in the *syn* conformation (so it was the 3+1 hybrid topology; Figure 4). The true extent of this phenomenon cannot be evaluated from the current simulations reliably because the εRMSD CV drives the RNA towards the all-*anti* parallel-stranded G4, i.e., the simulations are rather biased against the *syn* orientation of Gs in the later stages of the folding pathways. However, the *syn* G conformation is not prohibited and its occurrence in some of the misfolded rG4s indicates that structures with *syn* G orientation may be transiently populated during the folding attempts. It also indicates that folding of structures of rare rG4s with *syn*-oriented guanines^105, 106^ may arise by the same folding mechanism as the “common” all-*anti* rG4s.

The findings in this study thus offer the folding picture of the rG4 folding. At the atomistic level, it is a gradual, multi-pathway growth of the G4 from the coil-like ensemble via numerous diverse individual routes while avoiding sharply structured intermediates resembling the structure of the final rG4 (Figure 5, Supporting Figures S6 and S7). As noted above, the results may be obviously limited by the accuracy of the atomistic force field and the used collective variable. However, the inter-tract εRMSD CV should not prevent formation of rG4-like hairpin and triplex intermedites; in fact, it could rather support them. Obviously, due to the use of replica-exchange enhanced sampling methodology (among other reasons) the present simulations cannot provide any insights into the kinetics (duration) of the individual folding attempts and pathways. However, they should be capable to provide insights into the structural ensembles that are involved in the process. The suggested folding mechanism is consistent with the one predicted for the analogous DNA sequence.^66^ The structuring mechanism is also remarkably similar to the one theorized based on the simulation behavior of RNA G-hairpins and their preference of adopting the cross-like arrangement over the ideal G4-like parallel-stranded one.^32^ On the other hand, a recent coarse-grained (CG) simulation study suggested a different folding process.^77^ The authors initially observed formation of three-layered antiparallel G-hairpins and either two of these combined to form the rG4 directly, or a three-layered G-triplex occurred and then the rG4 was formed by attachment of the fourth strand (Figure 5). Thus, formation of columns of three stacked guanines was a key feature in the CG model, resembling simple models often used in indirect interpretation of primary experimental data. In our all-atom simulations, guanines tended to be stacked together, too, but structures with just two stacked guanines in a tract were more prominent, leading to the gradual formation of two-quartet rG4s intermediates. Such structures were absent in the CG-based model. Thus, although atomistic simulations and CG simulations studies both indicate a multi-pathway rG4 folding process, the nature of ‘multi-pathway’ differs markedly between the two (Figure 5). The CG model proposes a few routes with salient straightforward rG4-like intermediates, while the atomistic model suggests a multi-pathway process with numerous individual atomistic folding routes, starting from a coil-like ensemble, and largely avoiding well-defined rG4-like intermediates. The coil-like state can perhaps be likened to the molten globule state in the protein folding, with the stacked G-tracts resembling simple secondary structure elements. One reason for the difference is the nature of the CG model, which reduces a nucleotide into three beads and does not include the explicit solvent. This obviously leads to a less precise description of the interactions between nucleotides. It decreases the number of possible interactions and simplifies and smoothens the free-energy landscape, possibly pushing the system into regions that are structurally idealized. In other words, the CG model is unlikely to populate the coil-like ensemble. Our all-atom simulations, on the other hand, are affected by the used CV (inter-tract εRMSD), and one might argue that it likely pushes the guanines together to form the coil and might also promote formation of the two-quartet rG4 intermediate ensembles. We have tested alternative (and simpler) collective variables, namely the number of native H-bonds and the K^+^-coordination number, which in our opinion would have had smaller impact on affecting the native folding pathway(s). However, simulations using these CVs resulted neither in productive folding nor into structures at least remotely resembling partially-folded rG4 (data not shown), justifying the use of the inter-tract εRMSD metric. Clearly, without coarse-graining or enhanced-sampling with CV-based dimensionality reduction in all-atom MD simulations, spontaneous rG4 folding would be infeasible within the currently affordable simulation time scales. Each approach has limitations, and considering the potential impact of the methodological differences, it is possible that the actual folding mechanism may incorporate elements from both the CG model and the atomistic model presented here. We nevertheless think that the picture provided by the all-atom simulations is the one closer to reality, unaffected by simple assumption-based folding schemes found in the literature and often used in interpretation of experimental data.^22, 46^ Although the G4 molecules with their tetrads and bound ions are quite specific biomolecular structures, the G-rich nucleic acid chains should not fundamentally differ from the other biomolecular chains in their tendency to form compacted coil-like ensembles. The possible role of hidden (invisible) coil-like ensembles in G4 folding processes has been noted also in some experimental studies.^28,47^

**Figure 5.**
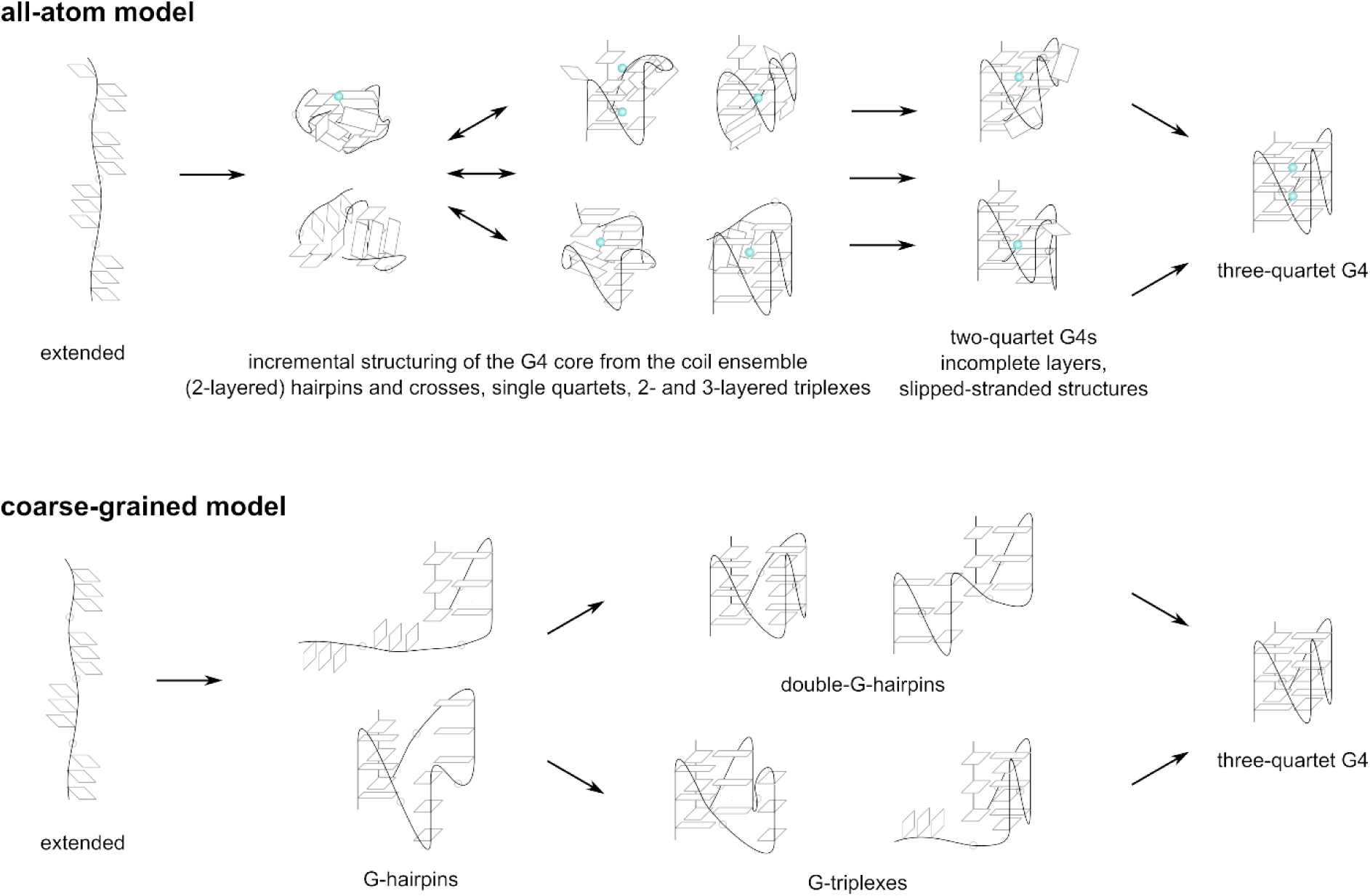
Generalized rG4 folding mechanism as suggested by all-atom simulations in the present work and previous CG simulations.^77^ The CG model proposes a multi-pathway folding mechanism involving a few well-defined G-hairpin-based intermediates, but does not specify when cations interact with the RNA. In contrast, the all-atom simulations reveal numerous atomistic pathways originating from a compact, unstructured coil-like ensemble, without the presence of well-ordered intermediates. Folding progresses incrementally, with structuring occurring within the coil-like state. During this process, cations are incorporated, and various two-quartet rG4 configurations emerge as important components of the late-stage folding transitory ensemble. Thus, while both models propose multi-pathway folding, their interpretations of the term differ significantly.

### RNA coil-like ensemble is looser than that of DNA

Radius of gyration (Rg) is sometimes used, in experimental as well as simulation studies, as a measure for determining whether G4 is folded or not. Small Rg values are implicitly assumed to indicate a folded state. However, Rg of the coil ensemble is smaller than that of the fully folded rG4 (calculated for the whole molecule; Figure 6), suggesting that Rg is not able to reliably discriminate between fully unfolded but compacted, partially unfolded and fully folded rG4 states. We have demonstrated this before also for dG4.^62, 66^ Interestingly, comparison of the values with our previous study on dG4^66^ reveals that while Rg of folded rG4 is comparable to dG4, Rg of the coil-like ensemble formed by DNA is smaller, suggesting the DNA ensemble is more compact than the RNA one.

**Figure 6.**
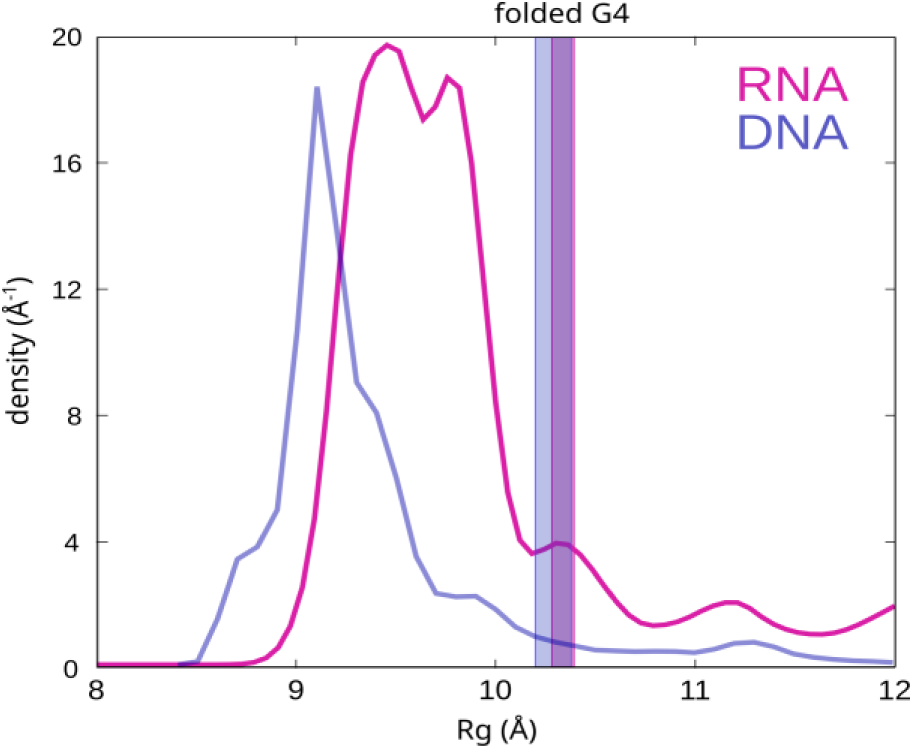
Reweighted Rg population distribution in ST-metaD reference replica of RNA simulation 1 (the one with the most folding events) and its comparison with distribution observed in a recent dG4 folding simulation.^66^ The vertical stripes correspond to the median Rg of the folded rG4 and dG4 ensemble, with a width of interquartile range.

### Limitation of the ST-metaD method with inter-tract εRMSD CV

ST-metaD simulations represent a high-end enhanced-sampling technique capable of overcoming free-energy barriers even in rather large systems, which was not computationally feasible until recently. A limitation that arises naturally from the metaD technique is its inability to accelerate sampling in directions orthogonal to the CV used. Identifying a single CV or a set of CVs that effectively capture the reaction coordinate (or, rather, mechanism) across a multidimensional free-energy space is often challenging. There is always a risk that important transitions may remain obscured by the chosen CVs. In cases where the conformational space is inherently highly multidimensional with many slow degrees of freedom, achieving a reliable dimensionality reduction may even be fundamentally impossible. We have previously suggested that, even though εRMSD is one of the best known CVs, especially the dG4 folding landscapes are so intrinsically multidimensional that their description is principally irreducible to one or few CVs, no matter how carefully chosen they are.^27, 74^ Therefore, we employed the replica-exchange solute tempering (ST) to help overcome enthalpic barriers in general and to somewhat alleviate the main drawback of metaD. However, the capabilities of the replica-exchange protocols to enhance sampling are also not unlimited.^107, 108^

Notably, another recently introduced approach for improving sampling along CVs is the On-the-fly Probability Enhanced Sampling method (OPES).^109^ It modifies the way CV is influenced by focusing on reconstructing the probability distribution rather than building a bias potential, as in metaD.^109^ Instead of depositing Gaussians to incrementally adjust the bias, OPES uses an on-the-fly kernel density estimate of the probability distribution to define the bias and has been suggested as a faster alternative to metaD for multidimensional free-energy surfaces where it is difficult to choose CVs.^109^ OPES has recently been used to explore the folding and conformational transitions between several dG4 topologies of the DNA human telomeric sequence.^58^ Although OPES can also be combined with parallel tempering methods,^110^ detailed testing involving transitions in complex systems is required to confirm its benefits over the pure metaD approach, particularly when combined with parallel tempering methods.

Even though the combined ST-metaD method sounds robust, we have encountered a rather undesired behavior in our simulations, which clearly illustrates limits of the enhanced sampling simulations. Extending the simulations beyond 2 µs did not necessarily yield more folding events as most folding events occurred in the early simulation stages (Figure 3). We think that this simulation development likely stems from a stronger drive toward the folded state that is applied at the simulation start in conjunction with a suboptimal CV; unfortunately, as mentioned above, it may actually not be possible to pick a CV that would capture the folding mechanism completely because of its intrinsic multidimensionality. In the later simulation stages after the rG4s was formed, we often observed only reversible unzipping of one or more guanines from the rG4 and in a few cases the rG4 was disrupted irreversibly. Once the bias starts reaching convergence in the second half of the simulations, the sampling of folding events becomes less efficient, i.e., the expected frequent reversible rG4 unfolding and refolding events did not happen. We admit that this limited structural sampling indeed may cast some doubts on the calculated rG4 ΔG_fold_ value, even though the bias was technically converged (Figure 2). We found out that also the previously reported simulations on dG4^66^ suffer from the same issue, so the relative comparison of the rG4 an dG4 ΔG_fold_ values could still be qualitatively correct. We also assume that the observed folding events are qualitatively representative from the structural point of view (Figures 4 and 5), considering the limitations that have already been discussed throughout the paper. Nevertheless, our observation is a reminder of the limitations that may be present even when using sophisticated enhanced-sampling methods like ST-metaD and which may not always be acknowledged in the literature. In fact, this issue might in principle be similar to the tradeoff between the convergence speed, number of folding events and the extent of exploration of the free-energy landscape caused by the bias deposition and suboptimal selection of CVs as described for the OPES method.^111^ Yet, even with this sampling issue, the ST-metaD is likely one of the best MD simulation techniques suited for studying the G4 folding problem at the all-atom resolution available to date. We would like to point out that no such problems were detected in our recent ST-metaD simulations of simple RNA stem-loop hairpins.^108, 112^ We plan to explore this issue in greater detail in future work.

## Conclusions

RNA G-quadruplexes (rG4s) have been recently identified as structural species likely involved in a variety of biological processes. Despite its importance, the folding mechanism of rG4s has not yet been fully understood at the atomistic level of description. In this work, we successfully folded the r(GGGA)_3_GGG sequence into the parallel-stranded rG4 using all-atom molecular dynamics (MD) simulations (Supporting Movies S1-S11). To achieve this goal, we employed the well-tempered metadynamics method coupled with solute tempering (ST-metaD), which enabled us to overcome the sampling limitations of standard unbiased MD simulations without the necessity to employ coarse-grained models. Our results suggest that the rG4-forming strand forms a compacted coil-like ensemble, in which consecutive guanines tend to stack to each other (Figure 3). The ensemble is very dynamical. The arising columns of two or more stacked guanines interact together and form various intermediate subensembles, such as cross-like triplexes, two-layered G-triplexes and most importantly two-quartet rG4s (Figure 4, Supporting Figures S6 and S7). The coil-like ensemble may thus functionally resemble the molten-globule state in protein folding. The monovalent ions extensively interact with the RNA (or DNA) chain already at this stage. Finally, the full three-quartet rG4 emerges from this transitory ensemble by step-by-step rearrangements involving incorporation of additional guanines, G-tract rotations, strand slippage movements and possibly some other transitions. The simulations also show that while RNA is capable of forming off-pathway misfolded rG4s to certain extent, their stability is rather low which highlights the key difference from dG4 folding landscapes.^27, 33, 74^ We thus demonstrate that the rG4 folding process is inherently multi-pathway. However, by ‘multi-pathway’ folding we mean a process where the final rG4 fold emerges from the compacted coil-like ensemble via numerous very diverse individual routes, and not a process that involves a few sharply defined intermediates possessing already the native Hoogsteen base pairing (Figure 5). Difference between the two diverse but at the same time complementary interpretations of the term ‘multi-pathway G4 folding’ are discussed in the paper.

As noted in the Introduction, earlier studies have attributed the long folding times of many DNA quadruplexes to kinetic partitioning.^21, 27, 28^ This phenomenon arises from the presence of multiple deep free-energy basins (long-lived folds, probably diverse quadruplexes) on the energy landscape, which compete with each other. The present results can be linked to this framework by proposing that also these competing dG4 folds emerge from the compacted coil-like ensemble, as visualized in this work. The kinetic partitioning for rG4 is largely eliminated since rG4 adopts exclusively all-*anti* parallel-stranded topology.^27,32,33^

While the simulations have provided mechanistic insights into the process of rG4 folding, they have also disclosed a severe force field imbalance. The calculated rG4 folding free energy is ∼ +12.5 kcal/mol, which is in striking disagreements with experiments. We assume that the prime reason of this discrepancy is the lack of polarization in the simulation force field leading to an imbalance between the folded and unfolded ensembles. We nevertheless suggest that the basic structural aspects of the atomistically-visualized folding events are relevant.

Although the composed ST-metaD enhanced sampling method has helped us to successfully overcome the high free-energy difference and positive free energy of the folded state (as predicted by the force field), we have faced issues intrinsic to the enhanced sampling technique itself and its use on a highly multidimensional free-energy surface. The ST-metaD trajectories experienced most of the folding events shortly after the simulation start, when the bias towards the target native rG4 was the strongest. In later stages the simulations became rather unproductive. This observation thus reminds us of limitations to consider and bear in mind when running enhanced sampling simulations of complex systems, which, in our opinion, are not always adequately acknowledged in the literature.

In summary, using enhanced-sampling all-atom MD simulations we have folded a parallel-stranded rG4 from a straight RNA chain, tracking down altogether eleven individual folding events in the continuous ST-metaD trajectories. Based on the simulations we suggest that rG4 folding is a multi-pathway process in which a compact coil-like ensemble plays a key role, forming a starting stage for the individual molecules to launch their folding attempts and structural transitions.

## Data Statement

Starting structures, reference structures, simulation protocols, and Supporting Movies are available at Github (github.com/ppokor/G4_folding_RNA) and simulation trajectories (reference replicas, reactive trajectories) and the calculated bias files are available on Zenodo (10.5281/zenodo.13646266).

## Funding

This work has been funded by the Czech Science Foundation grant number 23-05639S.

## Acknowledgments

The authors acknowledge Giovanni Bussi for fruitful discussion about the methodology limitations. This work has been conducted in the sustainability period of the project SYMBIT No. CZ.02.1.01/0.0/0.0/15_003/0000477 as its follow-up activity.

## Notes

### Competing Interest Statement

The authors have declared no competing interest.

