## Supplementary material for "RNA G-Quadruplex Emerges from a Coil-Like Ensemble via Multiple Pathways": S1 Supplementary Information

**Supporting Results**

In the following text, we provide description of the folding events observed in the four ST-metaD simulations. The reactive trajectories are labeled as given in main text Table 2 and the events and some of the structures are visualized in main text Figures 3 and 4, respectively. All the reactive trajectories are visualized in Supporting Movies S1-S11. The numbers in bold indicate which of the four G-tracts participate in a given structure, counting from the 5'-end.

<u>simulation #1:</u>

- <u>reactive trajectory 8</u> (1_1; reactive_traj_1_1.mp4):
  coil with crosses → 2-layered G4 → slip-stranded G4 → G4
- <u>reactive trajectory 10</u> (1_2; reactive_traj_1_2.mp4):
  2-layered hairpin →2-layered hairpin + one more G-tract to cross →2-layered triplex → G4 with G3 and G11 in the solvent (one tetrad in the middle and 2 triads) → G4
- <u>reactive trajectory 14</u> (1_3; reactive_traj_1_3.mp4):
  very loose 2-layered triplex →remaining G-tract to cross → 2-layered G4 → fluctuations in the outer layers → G4
- <u>reactive trajectory 7</u> (1_4; reactive_traj_1_4.mp4):
  hairpin → coil → **1-2-4** 2-layered triplex + G-tract **3** cross → **1-2-3** 2-layered triplex + G-tract **4** cross → G4
- <u>unproductive trajectory 1</u>:
  locked in 2-layered slip-stranded G4, G3 locked inside the loop

<u>simulation #2:</u>

- <u>reactive trajectory 5</u> (2_1; reactive_traj_2_1.mp4):
  U-shape with some pairs → circular with no specific pairs → coil → **1-4** 2-layer hairpin, G-tracts **2+3** cross → 2-layer G4 → G4 without one G → fluctuations with one G out and back → G4
- <u>reactive trajectory 6</u> (2_2; reactive_traj_2_2.mp4):
  coil with G-stacks →**3-4** 2-layer hairpin → one G from G-tract **3** coordinating Gs from G-tract **1** →unfold and back to coil with G-stacks →**1-4** 2-layer hairpin → **1-4** 3-layer hairpin, G-tract **3** interacts with it to do triplex but in antiparallel orientation→ G-tract **3** to cross and simultaneously **1-4** hairpin slip-strand, catching also one G from G-tract **2** by the G which is free to pair after the slip → G-tract **3** binds → 1-layer G4 → 2-layer triplex →3-layer triplex involving all 4 tracts → 2-layer G4 →G4
- <u>reactive trajectory 2</u> (2_3; reactive_traj_2_3.mp4):
  G-tract stacks → coil → **1-4** hairpin + G-tract **3** to cross → **1-3-4** triplex + G-tract **2** to cross → G4 without one G from G-tract **2** → G4

<u>simulation #3:</u>

- <u>reactive trajectory 4</u> (3_1; reactive_traj_3_1.mp4):
  **1-2** half-cross →cross-triplex **1-2-3** → 2-layered triplex →G4 with G15 stacked below G11 instead of being in plane with it → G4
- <u>reactive trajectory 10</u> (3_2; reactive_traj_3_2.mp4):
  **1-2** half-cross →**1-4** half-hairpin + G-tract **2** cross + G-tract **3** one G in resulting in 1-layered triplex → **1-3-4** 2-layered triplex + **2** cross → oscillating between 1- and 2-layered G4 with a triplex layer on top → locked in 2-layered G4 with first G11 out, then G3+G11 out, then G3 out →G3 stack with A and perpendicular to where it should be in G4 → G4
- <u>reactive trajectory 11</u> (3_3; reactive_traj_3_3.mp4):
  cross → hairpin → G-tract **3** cross to the hairpin→ 2-layered **1-2-3** triplex → slip-stranded G4 (slipped is the G-tract **4**, which was not in triplex) → G4

simulation #4:

- reactive trajectory 10 (4_1; reactive_traj_4_1.mp4):
  cross → hairpin → coil with 2-layred triplex and the remaining G-tract to cross → 2-layered G4 with strand-slip and G13 in the solvent → G4
- unproductive trajectory 6:
  locked in 2-layered G4 with G-tract **2** anti-parallel (G1 and G2 in *syn*, G3 forming the loop)

### Supporting Figures

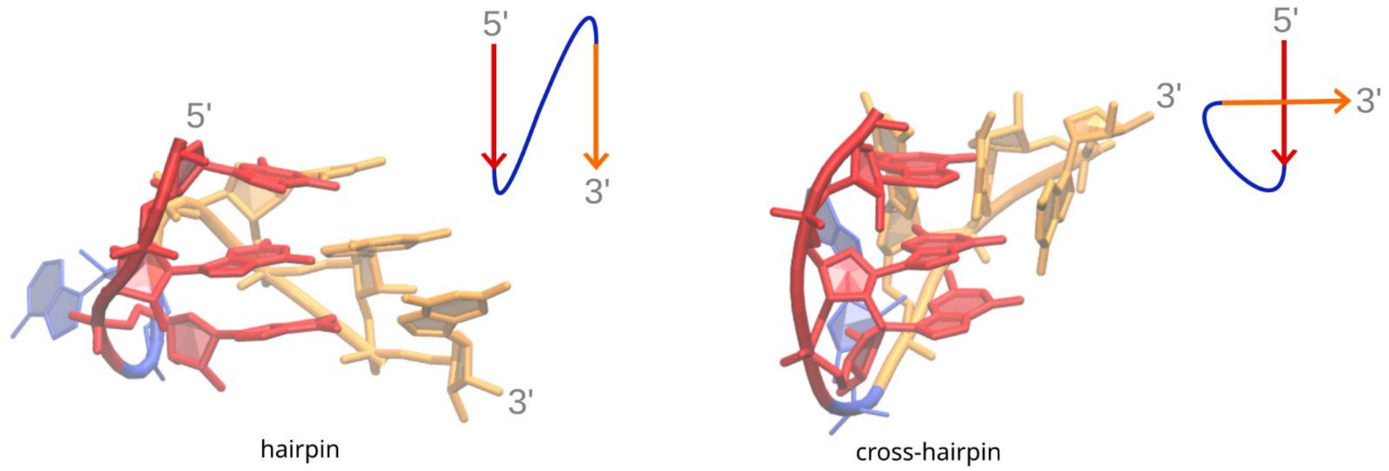


**Figure S1.** G4-like ("ideal") and cross-like hairpin. In the G4-like hairpin (G-hairpin), the guanines are H-bonded as in the native G4 structure, whereas in the cross-hairpin state (a quite broad ensemble of similar such states), the strands are mutually rotated without a single fixed H-bonding pattern. The same terminology applies to triplexes, i.e., G-triplex and cross-like triplex.

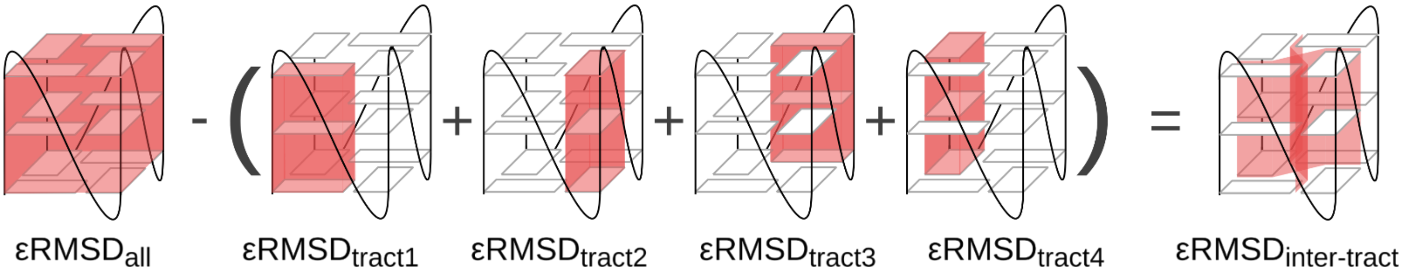


**Figure S2.** Schematic definition of inter-tract εRMSD for the full G4. εRMSD is a metric of structural similarity reflecting relative positions of nucleobases, including their interacting edges, designed specifically for nucleic acids.[1] The aim of the inter-tract εRMSD is to remove the contribution of intra-strand (i.e. within a G-tract) guanine stacking from the total rG4 εRMSD, thereby not promoting G-tract stacking by the simulation bias. In other words, the intrastrand (intra-tract) G stacking occurs spontaneously without the help of the CV-bias.

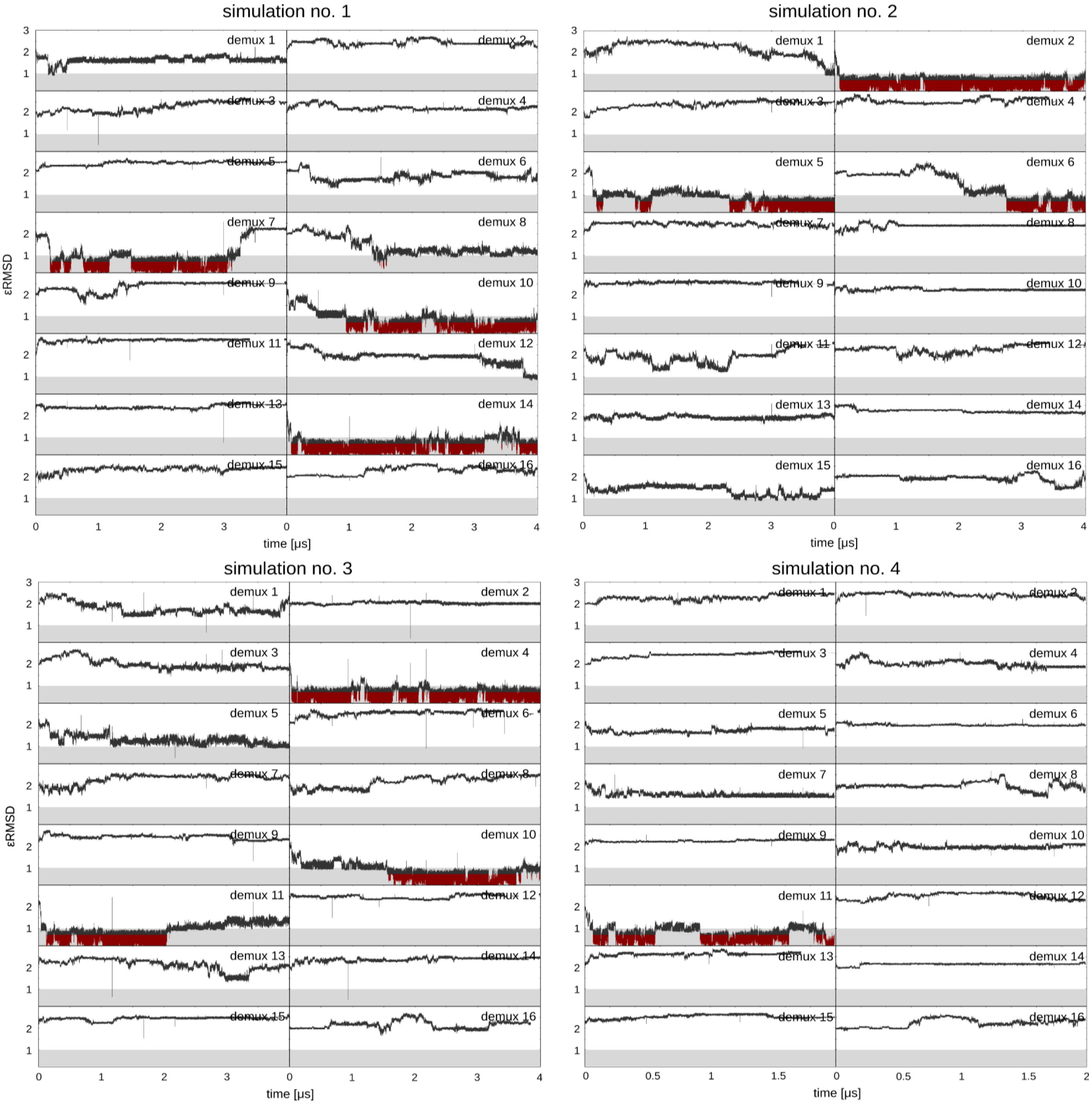


**Figure S3.** εRMSD development in the individual continuous (demuxed) trajectories. The grey area marks εRMSD < 1.0 while regions considered folded (εRMSD < 0.7) are marked with dark red. The development from the starting extended single strand state to the first event when it reaches εRMSD < 0.7 is considered a reactive trajectory. For example, in the simulation #1, demux 7 only the development until the first folding event is considered reactive trajectory. The refolding events, in which the already folded rG4 only partially unfolds and then refolds back are not considered proper folding events (e.g., simulation #1, demux 7, times around 0.9 and 1.6 μs). All the refolding events only sampled unstacking of guanines from the outer tetrads but no refolding passing through a loose coil state was observed. The graphs thus also show inefficient sampling of proper refolding once a fully folded rG4 is reached. This is an indication of a simulation limitation related to the ST-metaD technique and the chosen CV. This issue is in more detail discussed in the main text.

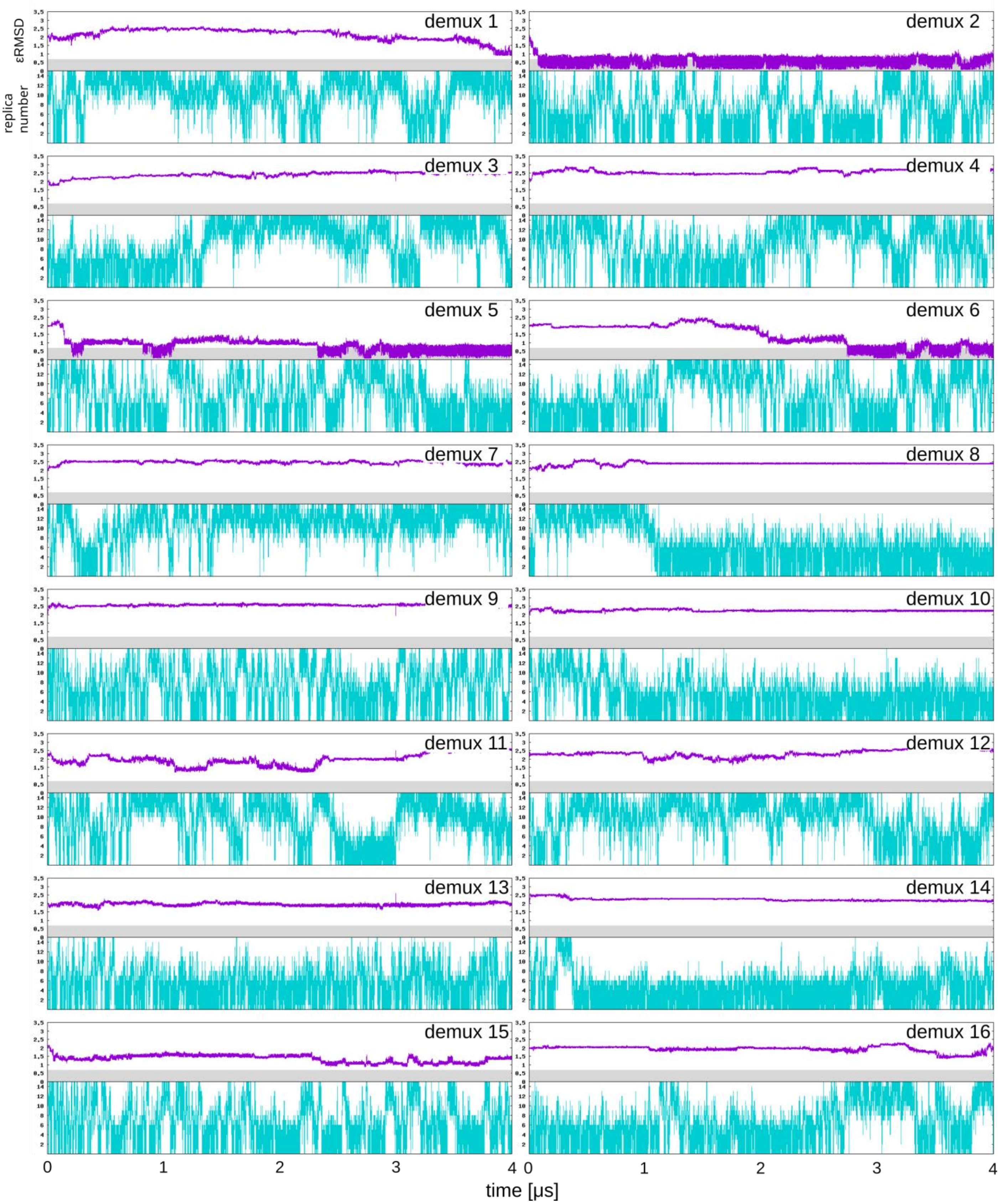


**Figure S4.** Individual continuous (demuxed) trajectories traveling through the space of 16 replicas of the REST2 ladder in simulation number 2. The lowest replica has the lowest effective temperature. The grey area in the εRMSD graph marked regions considered folded (εRMSD < 0.7).

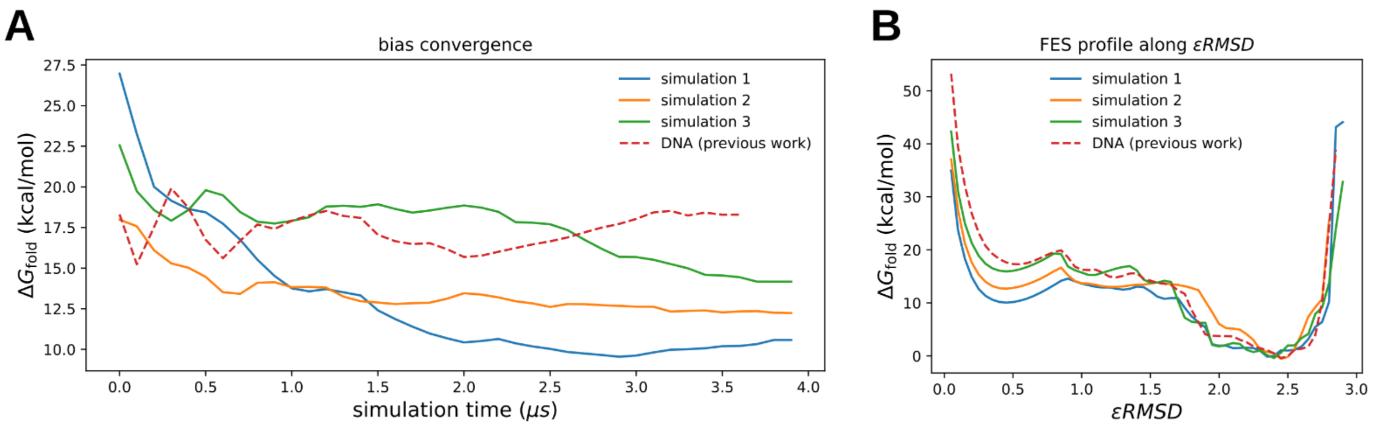


**Figure S5**. Comparison of folding free energy of rG4 and dG4 studied before.[2] (A) Bias convergence in the present RNA simulations and DNA (averaged data); the states with εRMSD to the reference G4 smaller than 0.7 were considered folded. (B) $\Delta G_{fold}$ along εRMSD, values are calculated using a bin width of 0.1. The shape of the curves is similar, with dG4 having the folded state ~4.5 kcal/mol higher than rG4.

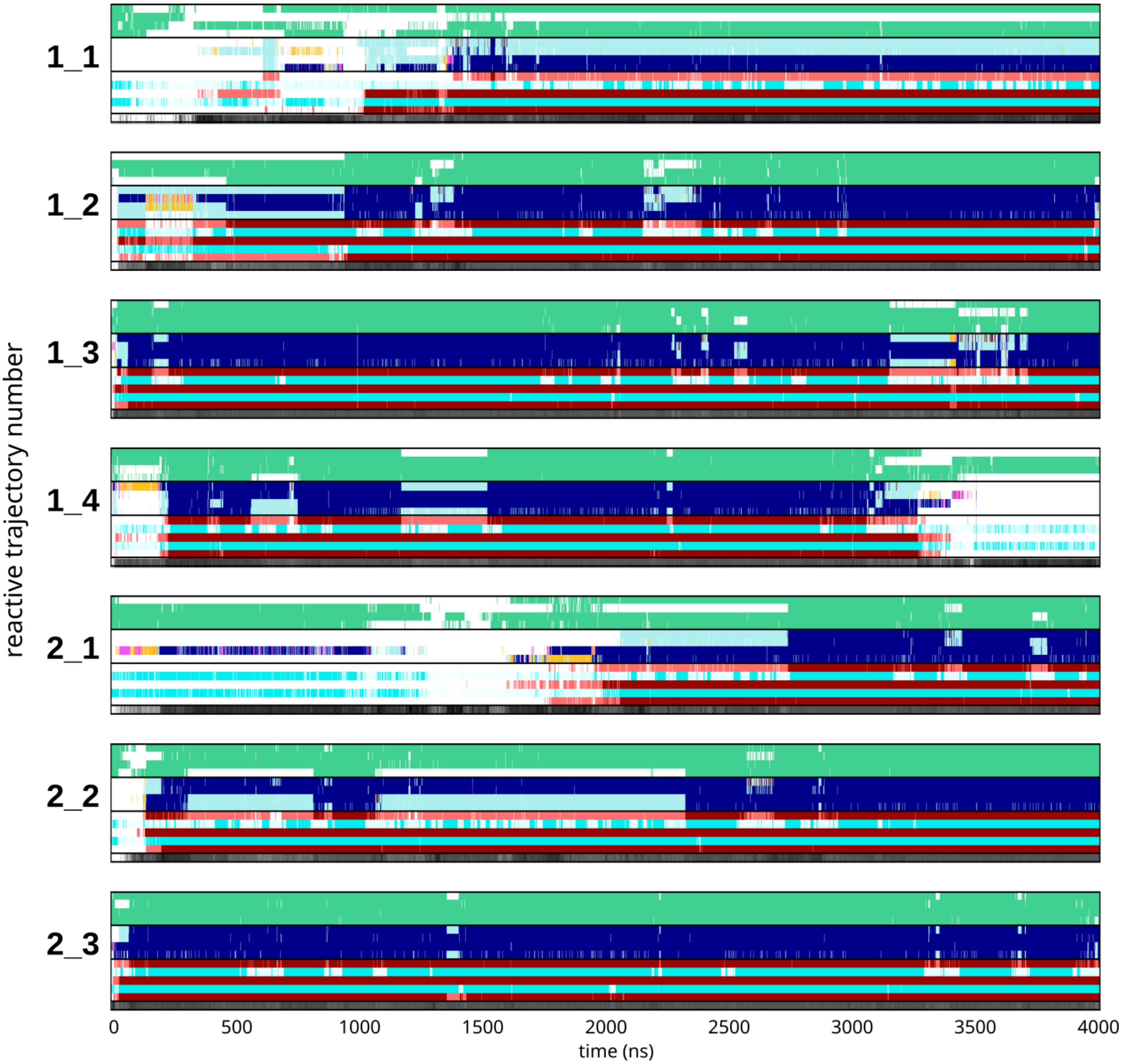


**Figure S6.** Development of key structural features in continuous trajectories in simulations #1 and #2 that contained rG4 folding event, i.e. they were productive. The respective reactive trajectories, i.e., the section that led from the unfolded state to the folded rG4 in a given trajectory, are shown in the main text Figure 3 while this figure shows the full 4 μs trajectories. For the legend, see Figure S7. Further details are given in the main text Table 2.

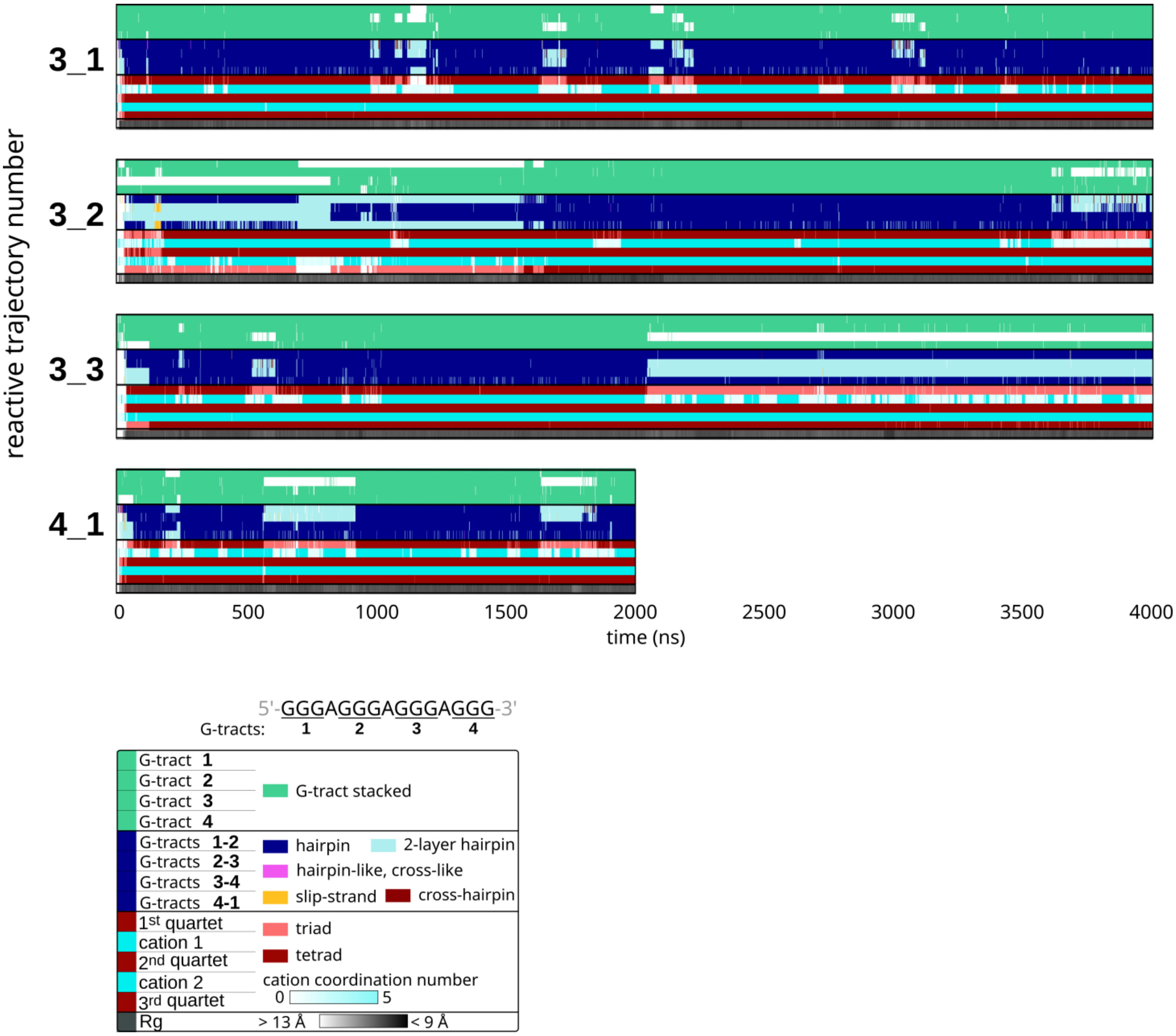


**Figure S7.** Development of key structural features in continuous trajectories in simulations #3 and #4 that contained rG4 folding event, i.e. they were productive. The respective reactive trajectories, i.e., the section that led from the unfolded state to the folded rG4 in a given trajectory, are shown in the main text Figure 3. Further details are given in the main text Table 2. The four top-most stripes in each graph monitor stacking of the G-tracts, the four stripes below monitor mutual orientations of the neighboring G-tracts, the next stripes indicate formation of individual G4 layers (triads or tetrads) and cation coordination between them, and the last stripe shows overall radius of gyration (Rg). When all the stripes are colored as in the legend, the G4 is fully formed.

### Supporting Movies

Reactive trajectories are visualized in Supporting Movies S1-S11. The order corresponds to the numbering of reactive trajectories (see the Supporting Results section above).
